# Movement Effort does not alter the Planning Horizon of Sequential Reaching Movements

**DOI:** 10.1101/2025.11.07.686755

**Authors:** Aslan Bellmann, Leonie Oostwoud Wijdenes, W. Pieter Medendorp

## Abstract

Everyday tasks often involve multiple reaching movements in a sequence, where the choices made for one movement also affect the conditions for subsequent movements. Previous research indicates that humans are capable of planning ahead to optimize such sequences. Here, we ask whether increasing the resistance to movement prompts participants to plan even further ahead to reduce overall movement effort. Participants (n=28) were shown 14 targets that varied in size, value and location, and were instructed to accumulate as many points as possible. To collect each target, they reached while holding the handle of a robotic manipandulum that applied a resistive force. We analyzed their target choices using a planning model that takes into account the size, value and distance of several future targets. We expected that, as resistance – and therefore the effort required to move between two targets – increased, participants would plan further ahead to minimize total movement distance. Results show that participants typically planned 2-4 targets ahead. However, increasing resistance did not significantly affect the planning horizon. We conclude that movement effort is not a primary constraint on the planning horizon in sequential reaching movements.

**NEW AND NOTEWORTHY:** Humans are capable of planning ahead in sequential reaching movements, but whether they adjust their planning horizon according to task demands is unknown. To manipulate task demands, we increased movement effort during a free target collection task. This makes planning more beneficial. However, we found no adjustment in planning horizon, suggesting that movement effort might not be a critical factor for planning in sequential reaching movements.

## 1 Introduction

Picture yourself seated at a breakfast table with friends. As one person requests the salt, another asks for the coffee can. Almost instinctively, you coordinate a sequence of movements to hand both items over without colliding with other objects on the table, all while reaching for the jam jar yourself. What mechanisms in the brain govern this sequential reach behaviour?

Compared to a single, discrete reach task, sequential reach tasks involve movements that are more continuous and blended, typically irregularly spaced and of varying amplitudes.

They may require rapid decisions about which target to acquire next, how to acquire it, and when to generate the response, ultimately leading to future states that allow subsequent action plans also to be feasible and relevant. Recent literature suggests that each movement may be prepared and executed in a way that takes into account subsequent movements—a process known as planning (Mattar and Lengyel 2022). Planning can improve movement optimization (Ariani et al. 2021; Bashford et al. 2022; Kashefi et al. 2024, 2025; Velázquez-Vargas and Taylor 2025), which typically entails balancing the minimization of effort and the maximization of reward (Carroll et al. 2019; Christopoulos and Schrater 2015; Pierrieau et al. 2021; Summerside et al. 2018).

Reseach indicates that individuals are capable of planning multiple movements ahead of their present action, and that this planning capability improves with experience (Ariani et al. 2021; Ariani and Diedrichsen 2019; Bashford et al. 2022; Diamond et al. 2017; Kalidindi and Crevecoeur 2024; Kashefi et al. 2024, 2025). When multiple targets are acquired in a fixed order, such as when typing in a phone number, the control of the current reach is tuned to subsequent reaches (Kalidindi and Crevecoeur 2024; Kashefi et al. 2024). Thus, future reaches are taken into account when choosing a control policy. Accordingly, if more information about future reaches is made available, movement execution time decreases and movement accuracy increases (Bashford et al. 2022; Kashefi et al. 2024, 2025).

When multiple targets are acquired without a fixed order, such as when wiping patches of ink off a whiteboard, research suggests that the selection of the next target is also influenced by the arrangement of the remaining targets. For example, Diamond et al. (2017) conducted a reaching study in which participants were tasked with collecting targets to score points. These targets varied in value, size, and location, and these properties were used to model participants’ choices. In this task, interdependence between reaches emerges because the choice of next target determines the distance to future targets, thereby influencing movement effort. The authors found that target choices were best explained by a model accounting for the next five movements. Notably, this planning effect was absent when participants performed the same task using eye movements, likely because target distance plays a lesser role in the effort required for eye movements. Hence, this suggests that planning in the human sensorimotor system arises when the interdependence between sequential movements sufficiently affects effort. This leads to the question whether the extent of planning can be modified accordingly. That is, if the interdependence of movements affects effort more strongly, do people plan even further ahead?

In the present study we further tested this hyposthesis by expanding upon the experimental paradigm developed by Diamond and colleagues. Participants were asked to collect targets while their movements were resisted using different viscous force fields. This modification was implemented based on the premise that as resistance to movement increases, the interdependence between reach movements becomes stronger. We hypothesized that this stronger interdependence would result in more extensitve planning.

## 2 Methods

### 2.1 Participants

Twenty-eight healthy participants (age 18-39 years, 24 female) with normal or corrected-to-normal vision and no known motor deficits took part in the experiment. Recruitment took place through a local recruitment platform. All participants gave informed consent before participating, and were reimbursed with 15EUR/h or an equivalent amount of course credits. The study was part of a research program approved by the ethics committee of the Faculty of Social Sciences, Radboud University, Netherlands.

### 2.2 Setup

Participants sat in front of a vBOT robotic manipandulum (Howard et al. 2009). They moved the vBOT handle in the horizontal plane using their dominant hand. Using a computer screen (ASUS VG278HR, 144Hz) and a mirror, we projected the virtual task environment onto the plane of movement. The handle position was sampled at 1kHz, and was displayed as a cursor (white circle with radius of 0.25cm). Participants had no direct view of the handle or their hand.

### 2.3 Task

In the experimental task, which we adapted from Diamond and colleagues (2017), a set of brown-colored targets was distributed across the workspace. Targets differed in location, size, and number of points they were worth (value). The objective was to score as many points as possible. Participants harvested points by moving the cursor into the target, upon which the target disappeared. Throughout the experiment, we imposed viscous force fields of varying strength, thereby altering the movement effort. An overview is shown in Figure 1.

**Figure 1.**
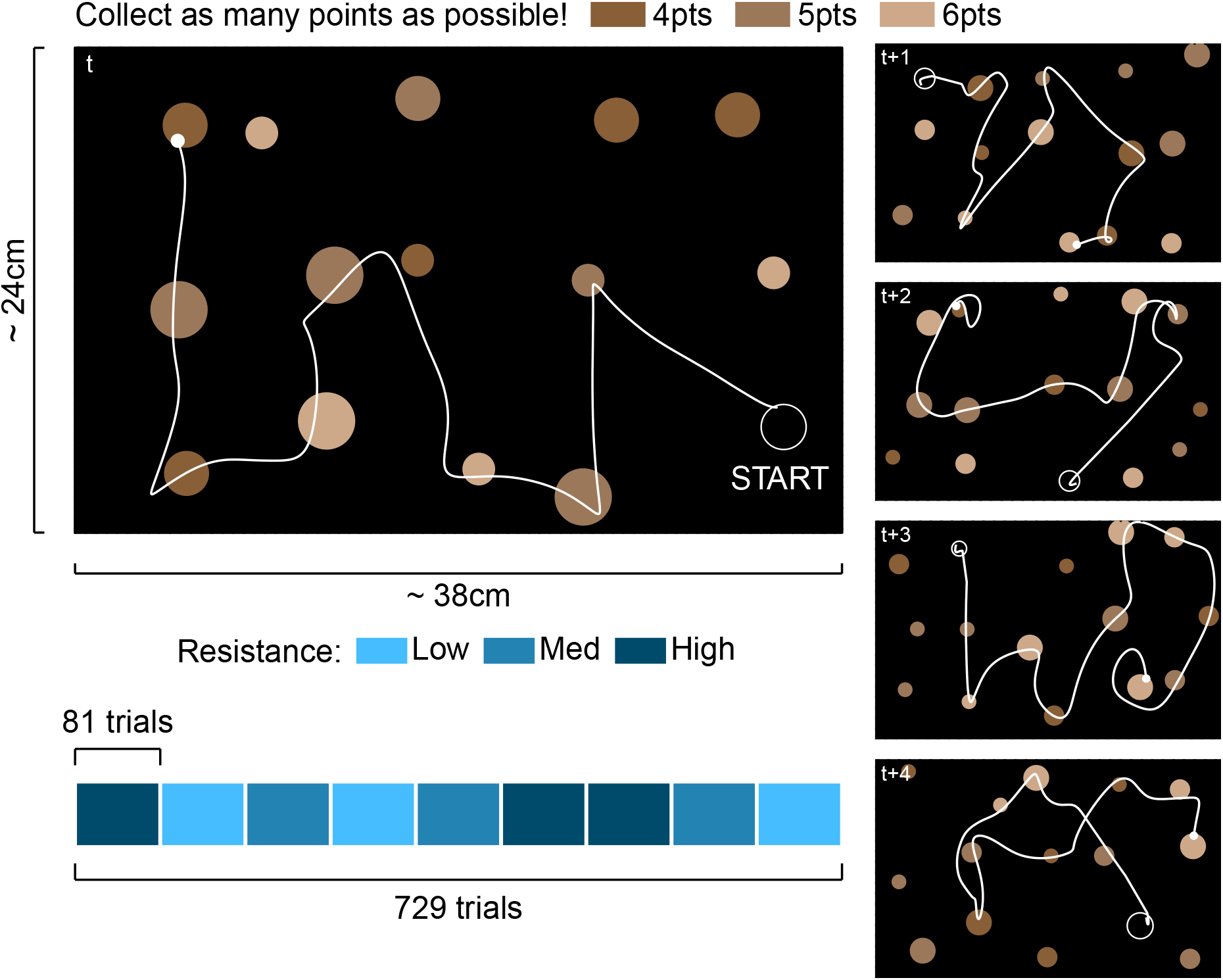
Task setup with example data from one participant. The white empty circle indicates the start postition for the trial (drawn here for clarity, but not visible during the trial). The white filled circle indicates the cursor at the final target collected during the trial. In this instance, a new set of targets appeared and the next trial started (see main text).

To construct the visual scene for a given trial, we divided the task workspace (37.5cm wide, 23.7cm tall) into an invisible 3×5 grid. We designed the task such that when a trial started, the cursor would always be in one of the 15 grid cells, while the other 14 grid cells would contain one target each. We computed target locations as the centers of the grid cells plus some uniform noise (4.7cm wide, 5.1cm tall; the maximum noise while avoiding overlapping targets). We randomly and equally assigned target radii of 0.8, 1.1, and 1.4cm and target values of 4, 5, and 6 points across grid cells. The value of a target was encoded by its brightness, with brighter targets being worth more points.

A trial always started with the cursor in one of the 15 grid cells and one target in each of the other 14 grid cells. Participants had between 2 and 3s to collect as many points as possible. The trial ended once a target was collected after the time had expired. Since the allowed time was random, it was impossible to predict with which collected target the trial would end and use that information to strategize. To make the task more engaging and to collect more data per unit time, we presented trials in sets of eight to ten. Within a set, trials followed each other continously: as soon as the final target was collected the workspace was wiped, 14 new targets appeared, and the next trial started immediately. Thus, the final collected target in trial *t* served as the starting position in trial *t* + 1. Between sets, participants could rest and initiate the next set of trials at their own pace, by placing the cursor within an indicated starting position for 2s. This starting position was randomized to one of the 15 grid cells.

After receiving task instructions, participants performed one practice block of 54 trials, followed by 9 experimental blocks of 81 trials each. Each experimental block had a resistive velocity-dependent force field with a strength of 0.01, 0.075, or 0.15 N/cm/s (henceforth simply referred to as low, medium and high resistance). The levels of resistance were randomly distributed across blocks, with the constraint that each level occurs once in the first, middle, and final three blocks. We did this to prevent any resistance-dependent effects to be confounded by time-dependent (e.g. learning) effects.

### 2.4 Model

To interpret target choices, we adapted the model proposed by Diamond and colleagues (2017). The model chooses the next target by jointly maximizing task performance and minimizing movement costs over the set of all future target sequences of length *h*.

Let *i* denote the last collected target and let *T* denote the set of targets that are still available, then the *step cost* from the last target *i* to the next target *j* ∈ *T* is

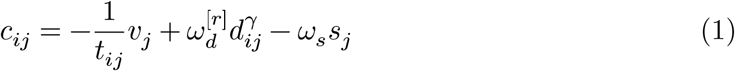

where *t*_*ij*_ is the expected movement time between targets, *v*_*j*_ is the value of the next target, *d*_*ij*_ the distance between targets, and *s*_*j*_ the size of the next target.

The first term in Equation 1 is the rate of reward, which reflects the task objective. The second term reflects that we expect movement cost to increase with distance. The free parameter *γ* ∈ (0, 5] allows for an exponential relationship. The third term reflects that movement cost may decrease with target size, as larger targets are easier to collect. The free parameters 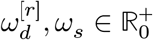 scale the contribution of distance and target size relative to the rate of reward. The superscript [*r*] indicates a free parameter that depends on the level of resistance.

For the expected movement time, we plugged in the median movement time, estimated via quantile regression conditioned on target distance and size. Movement times were positively skewed, with the longest times mostly due to targets that were missed initially. We chose the median, as it is less affected by extreme values.

The model chooses the next target *j* based on the best sequence of *h* ∈ {1, 2, 3, 4, 5} targets. Let *j, k, l, m, n* ∈ *T* be distinct available targets, then the *planning cost* of choosing target *j* is

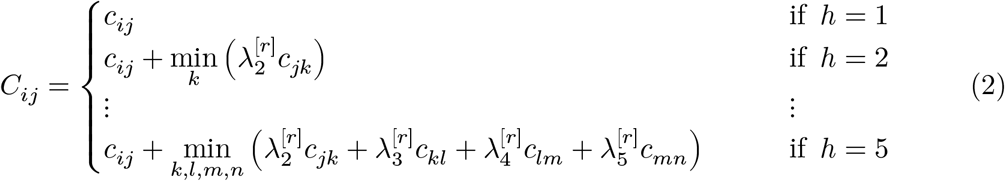

where the planning horizon *h* indicates how many steps are considered, and the free parameters 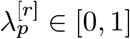 with *p* = 2, …, *h* reflect the importance of future steps. Note that if the planning horizon exceeds the number of available targets left, that is *h* > |*T* |, then the equation corresponding to *h* = |*T* | is used instead.

We verified that a greater planning horizon (*h*) leads to a sequence of target choices with a lower total cost (i.e. a lower sum of the step costs in Equation 1) via simulations shown in Figure S1. For intuition, consider that at each step, there is a configuration of targets and an associated distribution of available step costs. Without planning, the next target is chosen based on its step cost alone. With planning, the next target is chosen not only based on its step cost, but also based on subsequent target configurations. Thus, the chosen target in itself is less optimal (a higher step cost is chosen from the available distribution), however, this is masked by the target configurations being more favorable (the distribution of available step costs is lower). Consequently, the net effect is a decrease in total cost. Eventually, after most of the targets have been collected, only less favorable target configurations are left and planning becomes less beneficial (see gray segments in Figure S1). Note that in our task, it was almost impossible to reach this stage due to the limited time available for target collection, so planning would be beneficial (in principle).

We limited our analysis to a planning horizon of five steps maximum. First, as sequence length increases, combinatorial complexity makes model fitting infeasible. Second, simulations showed that increasing the planning horizon leads to diminishing decreases in cost (see Figure S1), thus the benefit of larger planning horizons is doubtful.

To account for perceptual noise, decision making noise, and motor execution noise, the ultimate target choice is treated as a probabillistic process. Given the previous target *i*, the probability of choosing target *j* is determined using the softmax function

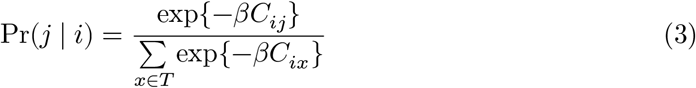

where 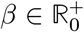 is a free parameter. Given our task has three resistance levels, the model has 3 + 3*h* free parameters: 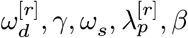. For example, if *h* = 2, the model has 9 free parameters: 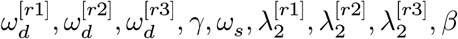.

### 2.5 Model Fitting and Comparison

For each participant, we fitted the above model for each value of *h* ∈ {1, 2, 3, 4, 5}. We excluded the first target collected on every trial, as the continous nature of the task meant that the choice of first target tended to be strongly influenced by the previous trial’s target configuration. We used the L-BFGS-B optimization algorithm (Byrd et al. 1995) to estimate the parameters that minimize the negative log likelihood. For each value of *h*, we initialized the optimization from various initial parameter sets (18 different initial guesses for *h* = 1, and 36 initial guesses for each *h* > 1 due to the additional parameters 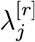) and selected the final parameter set yielding the lowest negative log likelihood. To compare models aross different values of *h*, we used the Bayesian Information Criterion (BIC)

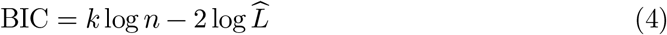

where 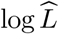 is the maximized log likelihood, *n* is the number of collected targets throughout the task, and *k* is the number of free parameters.

### 2.6 Statistical Analysis

More extensive planning can lead to a larger planning horizon (*h*) and/or to higher weights for future targets (*λ*_2_, …, *λ*_*h*_), suggesting that future targets exert a greater influence on the choice of next target. To compare planning horizons across resistance levels, we used the Friedman test. We opted for this rank-based test because the planning horizon values are discrete. To compare the weights 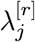 for the *j*th target ahead, we used a linear mixed effects model of the form `y ∼ j + r + (1|id)` where `j` is the target index, `r` is the resistance level, and `(1|id)` specifies a random intercept for each participant. The modelling approach accommodates cases where not every participant has weights for each target index due to different planning horizons. When computing pairwise contrasts from the mixed model, we adjusted *p*-values for multiple comparisons using Tukey’s method. Contrasts were considered statistically significant if *p* < 0.05.

### 2.7 Software

The task was programmed in MATLAB R2023b (The MathWorks Inc. 2023). Data processing, analysis, and visualization was done using code written in R 4.4.3 (R Core Team 2025), with the R-packages `targets` (Landau et al. 2025b), `crew` (Landau et al. 2025c) and `crew.cluster` (Landau et al. 2025a) being central to the pipeline. Dependencies were managed using the Nix package manager (Dolstra 2006) or the R-package `renv` (Ushey and Wickham 2025). The code will be made openly available at the Radboud Data Repository.

## 3 Results

Figure 2 illustrates overall task performance by displaying each participant’s cumulative score, as well as the number of points and the number of targets collected per trial. Although all participants successfully completed the task, their performance varied considerably. Across the entire experiment, participants scored between 35% and 56% of the total available points (Figure 2A). Higher scores reflected more points collected in individual trials, with the trial-to-trial variability similar across participants (Figure 2B). Higher scores generally corresponded with more targets collected. However, some participants prioritized collecting a greater number of targets while others focused on acquiring high value targets (Figure 2C).

**Figure 2.**
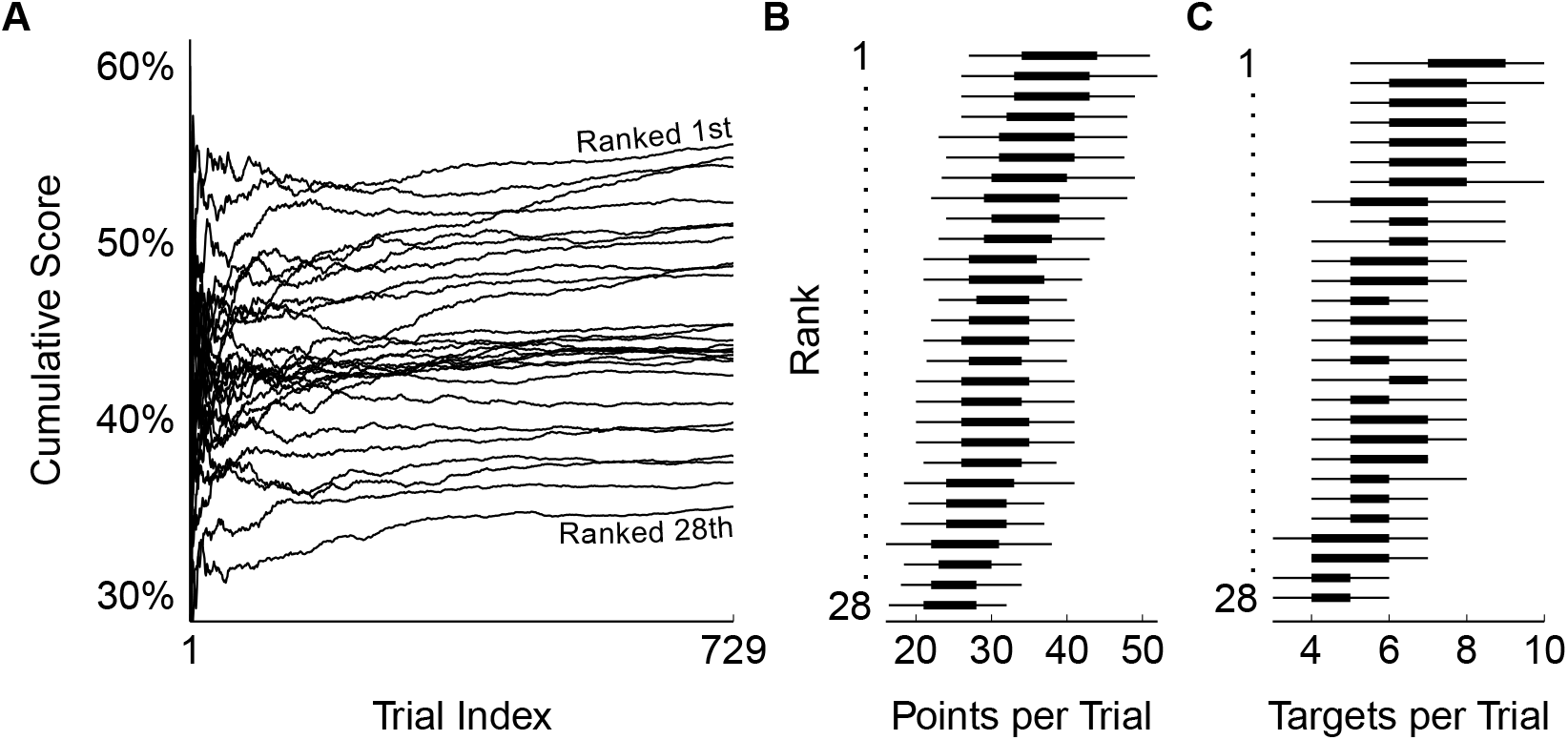
Overall task performance. **A:** Cumulative percentage of points collected vs. trial number. Participants are ranked according to the final percentage of points; this rank is used for the y-axis in panels B and C. **B:** Number of points collected per trial (maximum possible: 54-56 points) in ranked participant order. Intervals represent the middle 50% (bold) and 90% (solide line) ranges. **C:** Number of targets collected per trial (maximum possible: 14 targets). Intervals represent the middle 50% (bold) and 90% (solid lines) ranges.

Throughout the task, participants continuously had to decide which target to acquire next, as time constraints made it impossible to collect all of them. Figure 3 shows the harvesting preferences for the three manipulated target properties – distance, size, and value – by presenting, for each move, the normalized log ratio of the probability distributions for chosen targets compared to all available targets, separately for the three resistance levels. As expected, participants preferred targets that were closer, larger, and more valuable.

**Figure 3.**
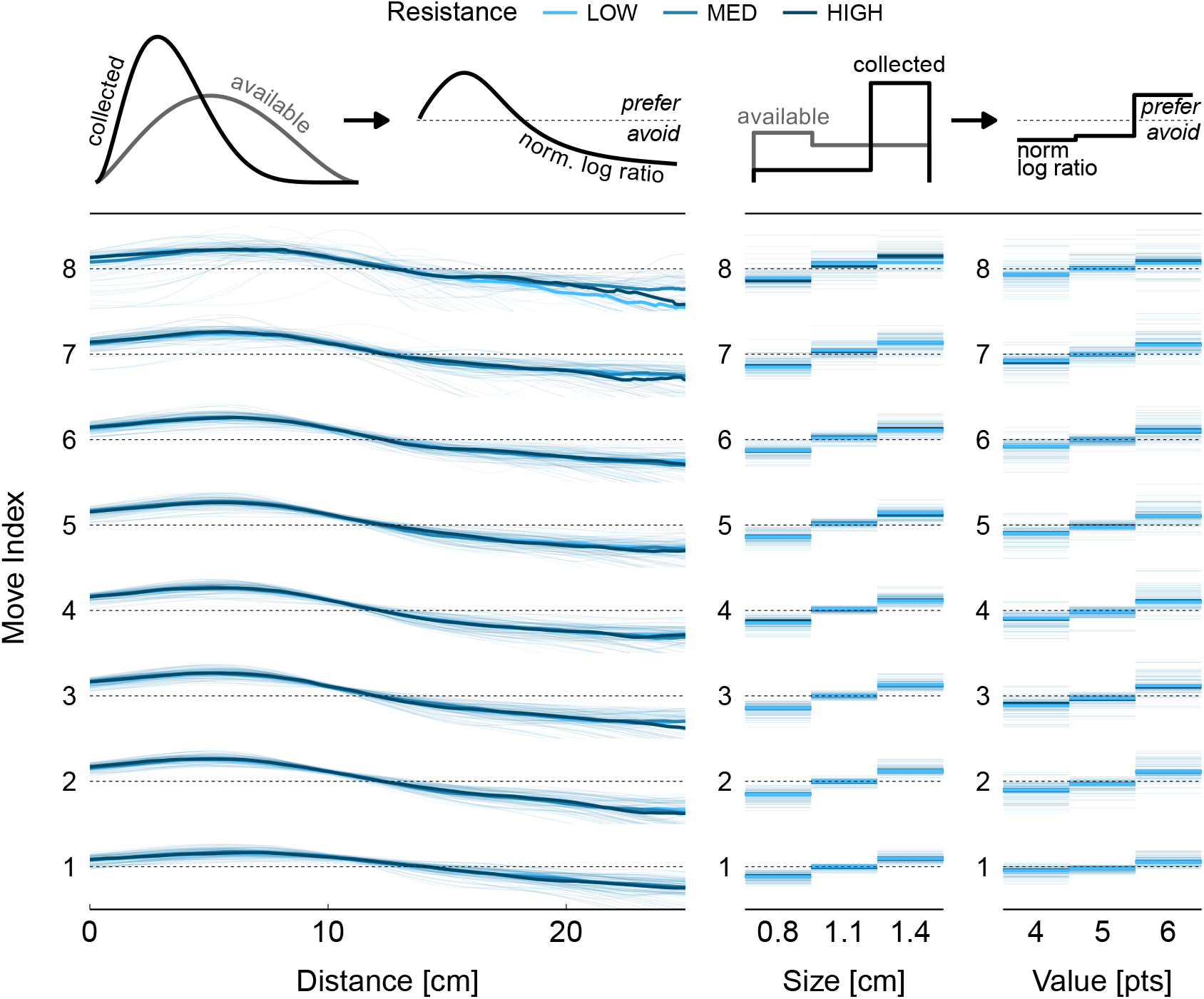
Normalized log ratio of probability distributions for chosen vs. available targets, for each target property. In mathematical terms, 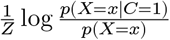 with the normalization factor *Z* = − log *p*(*X* = *x, C* = 1). This is equivalent to the normalized pointwise mutual information between a specific target distance, size, or value (*X* = *x*) and the decision to collect the target (*C* = 1). The dashed line corresponds to a value of zero. Values above the dashed line indicate a preference (i.e. targets with these properties were chosen more often than if chosen at random); values below the dashed line indicate avoidance. Bold lines are the group median, thin lines are individual participants.

We hypothesized that greater resistance to movement would prompt more extensive planning. Specifically, we expected that as resistance level increases, participant’s first moves would cover a longer distance to position the hand in a more advantageous area of the workspace for subsequent moves. However, Figure 3 shows that preferred target distances were consistent across resistance levels, indicating that the extent of planning did not vary. Similarly, the patterns of preference for target size and target value were consistent across resistance levels. Hence, resistance level did not appear to affect group-level strategies aimed at collecting more targets or maximizing value.

To quantify the degree of planning, we fitted models with different planning horizons (*h*) to the data, where the next target is selected by considering the next *h* ∈ {1, 2, …, 5} potential targets. If planning increases with resistance level, we would expect that models with larger *h* values provide a better fit to the data. Figure 4B shows the BIC score as a function of the planning horizon, normalized to the mean of the model with planning horizon *h* = 1. For most participants, increasing the planning horizon resulted in a lower BIC score (indicating a better fit), with the most pronounced improvements up to a planning horizon of 3 steps.

**Figure 4.**
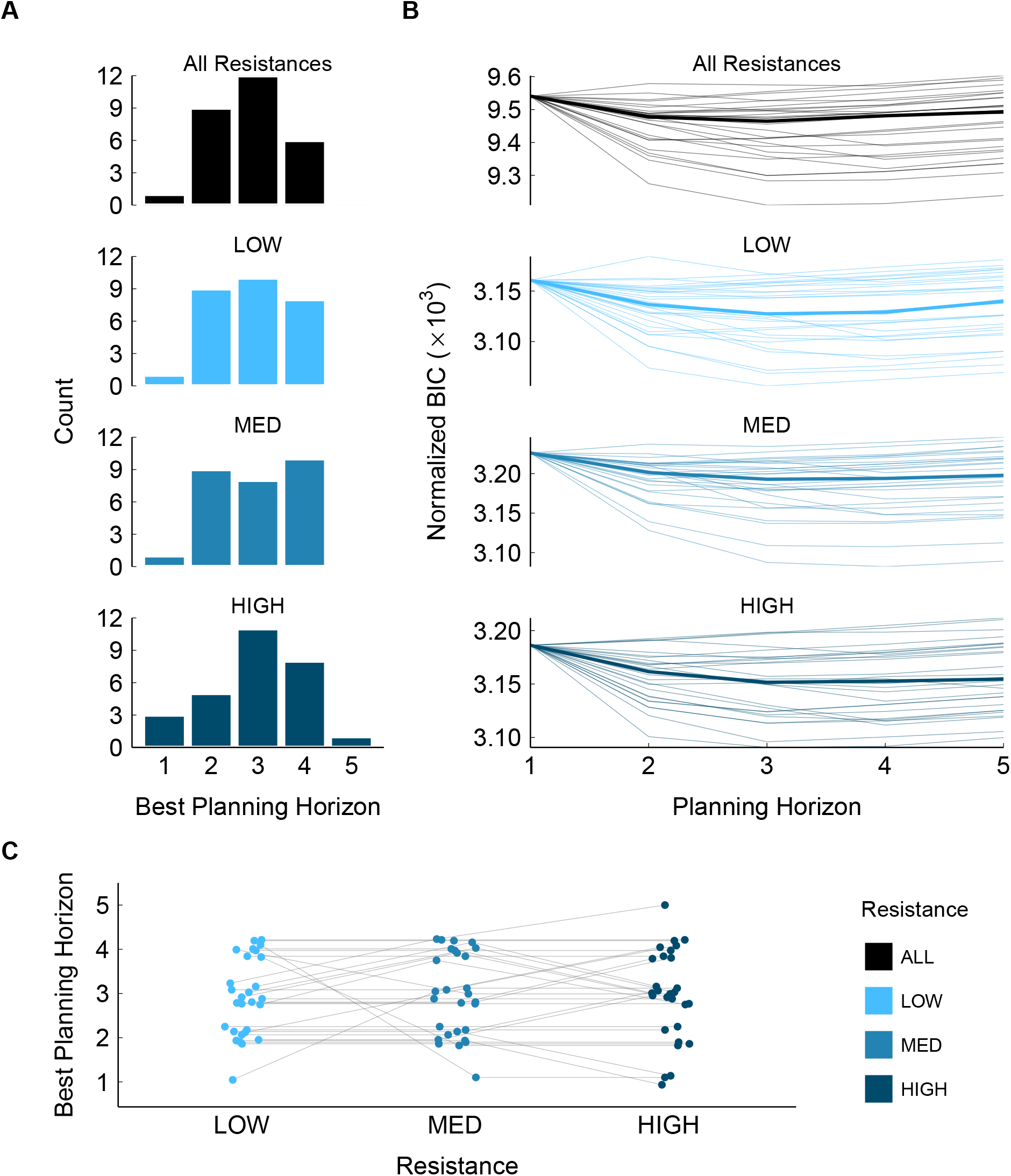
Model comparison. **A:** Histogram of number of participants whose data was best explained by the model with a given planning horizon. **B:** BIC for models with different planning horizons, normalized to the mean BIC of the model with planning horizon of 1. Bold lines show the group median, thin lines show individual participants. **C:** Within-participant change of the planning horizon best exlaining the data, as a function of resistance level. Data points and their connecting lines show individual participants.

Figure 4A presents histograms showing how many participants were best fit by models with a planning horizon between 1 and 5, seperately for each resistance level as well as aggregated across all resistance levels. The best fitting model typically had a planning horizon between between 2 and 4 steps. Figure 4C shows the best-fitting planning horizon plotted against resistance level, with lines connecting data points for individual participants. There was no systematic change in the best-fitting planning horizon as resistance increased. This was supported statistically: a Friedman test revealed no significant effect of resistance level on planning horizon size (*p* = 0.6).

While our analysis to this point suggests that our resistance manipulation does not significantly impact the planning horizon (*h*), it remains possible that it influences how participants weigh the importance of upcoming targets, as captured by the weights *λ*_2_, …, *λ*_*h*_. We show the weights from the best fit models in Figure 5. This reveals a subtle trend: as targets are further in the future, their associated weights gradually decrease, suggesting that nearer-term targets are prioritized. However, there was no effect of resistance level (all *p* > 0.05; the corresponding statistical contrasts are shown in Table S1 and Table S2).

**Figure 5.**
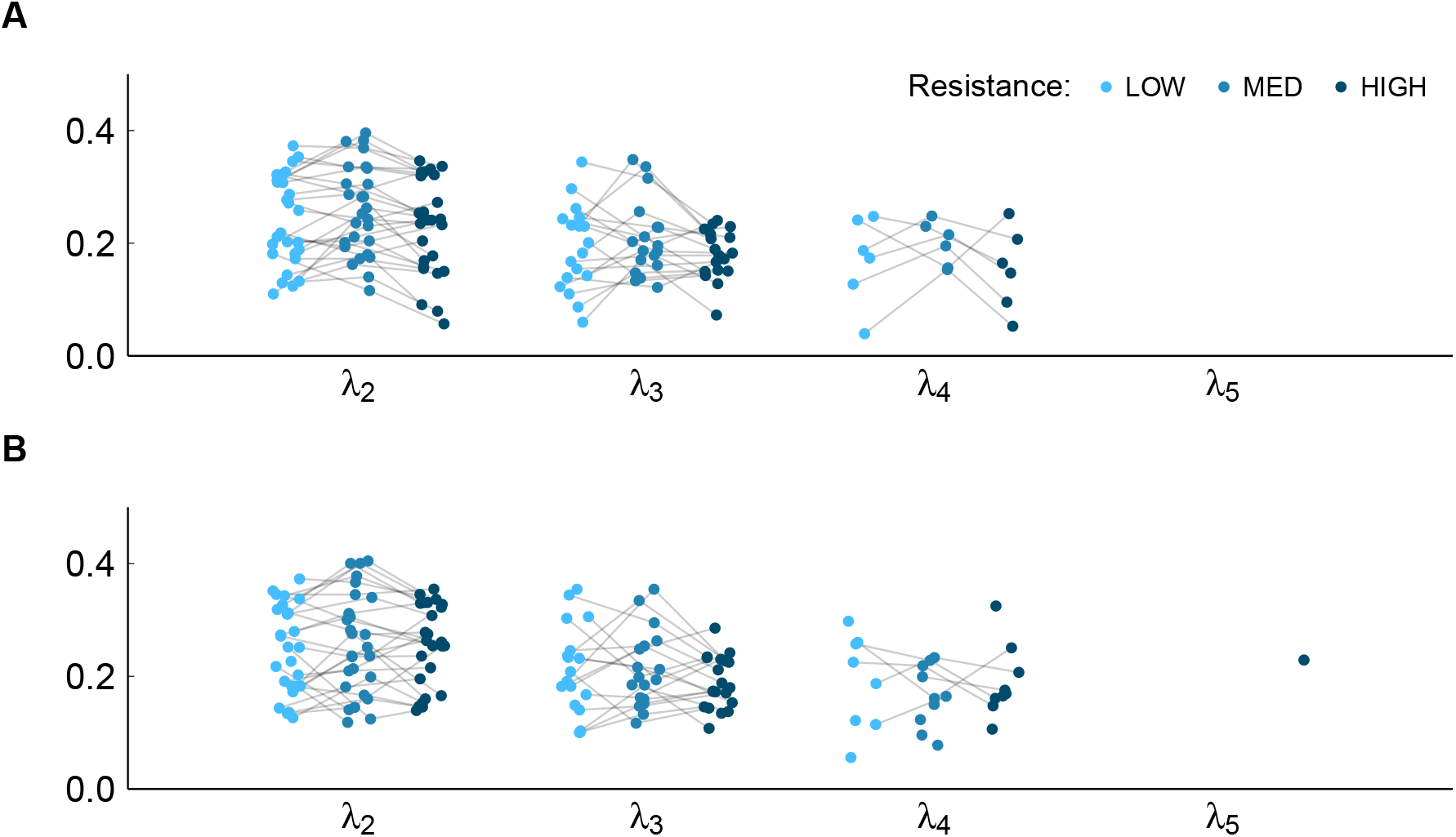
Within-participant change of the weights for future targets, between low (light blue), medium (blue) and high (dark blue) resistance. **A:** For a given participant, all parameters are taken from the model best explaining the data across all resistance levels. **B:** For a given participant, the parameters for a given resistance level are taken from the model best explaining that resistance level.

Finally, to assess whether our planning model accurately captures participant behaviour, Figure 6 shows how well the model predicted the targets chosen by the participants. Across all resistance levels, the models consistently predicted the chosen targets with an accuracy well above chance (Figure 6A). Notably, the targets selected by participants were generally among the three most probable targets according to the model (Figure 6B). Conversely, targets that were not chosen by the participants were rarely assigned high probabilities by the model (Figure 6C), further supporting the model’s predicitve validity. In summary, these results provide strong evidence that the fitted planning model offers a good description of our participants’ reaching behaviour.

**Figure 6.**
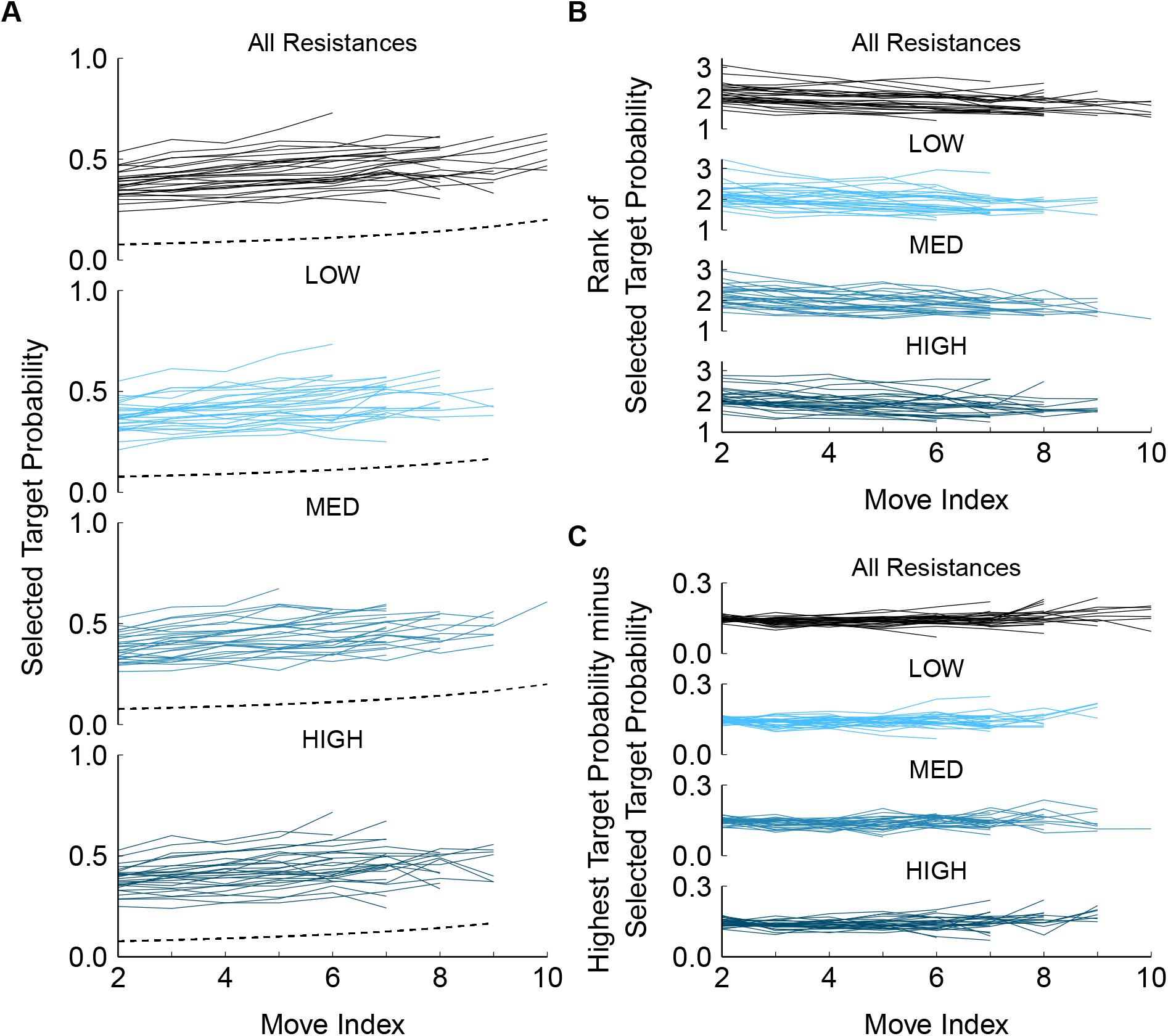
Predictions of participant behaviour according to the best fit model. For each participant, move indices for which at least 30 samples were available are shown. **A:** Average probability of the targets selected by participants, according to the best-fitting model. Dashed line indicates the probability if the targets were chosen at random. **B:** Average rank of the target selected by participants, with rank one corresponding to the most probable target. **C:** Probability of the best targets relative to the targets selected by the participants.

## 4 Discussion

Real world tasks often involve multiple reaching movements in sequence, where decisions about the next movement affect the options available for subsequent ones. Humans have been shown to leverage such interdependence by planning ahead multiple movements. Consequently, we hypothesized that increasing this interdependence would result in an even larger planning horizon. To test this, we used a reaching task where participants had to collect targets which were distributed across the workspace, and we modelled their choices as a planning process. We found that participants typically planned 2-4 steps ahead, that is, they chose the next target by also considering which future targets this would make available. We hypothesised that increasing the movement effort via a resistive force would prompt more extensive planning, as increased interdependence between movements makes planning more beneficial. However, based on model fitting, participants did not alter their planning horizon for higher levels of resistance. Thus, while participants used planning as an integral part of their decision making process, the extent of planning was not dynamically modified by movement effort.

In our task, interdependence between movements arises because the choice of next target determines the distances to subsequent targets. Participants could therefore optimize their choices by planning ahead: choosing targets that are not only desirable themselves, but also have other desirable targets in close vicinity. Simulations show that such planning becomes more beneficial as the importance of the distance between targets increases (Figure S1). Yet, a higher resistance to movement—which makes moving a given distance more effortful—did not increase the planning horizon in our study.

Our results suggest that movement effort is simply not an important decision making variable in reaching movements, at least given the levels of effort required in our task. In support, when reaching movements are modelled as optimal feedback control, there is evidence that kinematic (and not dynamic) variables dominate the cost function that is being minimized (Kistemaker et al. 2010, 2014; Mistry et al. 2013). Furthermore, in perceptuomotor tasks involving reaching movements, movement effort does not affect decision making while reaching distance does (Assarioti et al. 2025; Burk et al. 2014; Manzone and Welsh 2023). In particular, Burk et al. (2014) failed to see effects of resistive velocity-dependent force fields of 0.9 N/cm/s — considerably higher than our maximum resistance of 0.15 N/cm/s. Although their force fields were turned on sporadically while ours were on continously, this might indicate that much higher resistances are needed before the increase in movement effort affects decision making. This is also consistent with the fact that the weight parameter for distance 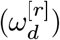 did not systematically increase with resistance level (Figure S2). To summarise, manipulations that involve kinematics or stronger forces may be needed to prompt a change in planning horizon.

Our finding that participants planned between 2-4 steps ahead aligns with previous research on sequential reaching movements (Kashefi et al. 2024), finger presses (Ariani et al. 2021; Ariani and Diedrichsen 2019), and combined motor and cognitive tasks (Velázquez-Vargas and Taylor 2025). However, it is somewhat in conflict with the study by Diamond and colleagues, which formed the basis for the present study. Using a similar reaching task and the same model, they found that most participants planned 5 steps ahead (Diamond et al. 2017). We can only speculate as to the cause for this difference, but it is worth highlighting some differences between the tasks. First, in the task by Diamond and colleagues, to collect a target, participants not only had to move the cursor into position, but also press a button. The need to coordinate these two actions may induce different control demands such as slowing down more at the target position, in which case distance may become a more important variable due to Fitts’ law (Fitts 1954), thereby prompting more planning. Second, their workspace (30 × 18 cm) was slightly smaller than ours (38 × 24 cm). It is possible that participants were better able to visually sample their whole workspace, thereby having more complete information available for their planning process. This information must predominantly come from peripheral vision, as humans tend to fixate a target until it is acquired, with the exact fixation point being skewed towards subsequent planned movements (Eluchans et al. 2024; Kashefi et al. 2024). Finally, differences may arise during model fitting. In particular, when finding the parameters that minimize the negative log likelihood, there is no guarantee to find a global minimum as optimization algorithms may get stuck in different local minima, depending on the initial parameter guesses. While we tried to mitigate this risk by initializing the optimization algorithm from many initial guesses, it remains a source of uncertainty. Overall, humans appear to plan around 2-4 steps ahead in sequential movements; how sensitive this planning horizon is to task demands remains a question to be answered.

In our task, whether a target has been selected in isolation or by planning ahead is not visible from behavioural data alone. Therefore, we used a model to assess the extent to which target selection was influenced by subsequent target configurations. We view this as a statistical (“what”) model, but it can serve as a starting point to discuss questions that arise from a mechanistic (“how”) viewpoint. For example, our model assumes that the costs for future targets, which are needed for planning, are known. One question is how these costs are represented in the brain. Goal-directed reaching movements, viewed through the lens of optimal feedback control, are determined by minimizing a cost function that at least partly depends on the properties of the target. If multiple targets are available, the cost function is minimized for each one, such that the target with the lowest cost can be selected (De Comite et al. 2023; Nashed et al. 2014). If planning relied on the same principle, it would require the minimization of an increasingly large number of cost functions—which seems infeasible, especially with limited time. A more efficient way would be to use heuristic information for planning, analogous to how some algorithms solve path finding problems (Hart et al. 1968). Our model also asserts that information about future targets is used to select a single next target, but in principle, a sequence of next targets could be selected at once (if a good sequence is available). With the sequence of targets known, reach control can be optimized across the sequence as a whole (Kalidindi and Crevecoeur 2024; Kashefi et al. 2024). This could be implemented as a hierarchical process: once a sequence of certain size is selected, higher level parietal and premotor area activity can represent the properties of that sequence and orchestrate lower level primary motor cortical activity representing individual movements (Ariani et al. 2025; Berlot et al. 2021; Gallivan et al. 2016; Yokoi and Diedrichsen 2019; Zimnik and Churchland 2021). Further behavioural modelling and neuroimaging studies will be required to unveil the mechanisms underlying planning in fast sequential reaching movements.

In summary, congruent with previous findings, we have shown that human decision making in a sequential reaching task is best described as a planning process. That is, humans leverage the interdependence between sequential reaches by accounting for future options in present choices. Our findings indicate that movement effort may not be a fundamental component of this planning process, thereby imposing additional constraints on the search for its neural correlates.

## 5 Data Availability

Source data for this study will be made openly available at the Radboud Data Repository.

## 6 Acknowledgements

We wish to thank Luc Selen for assisting with the technical setup of the experiment.

## 9 Supplemental Material

**Figure S1.**
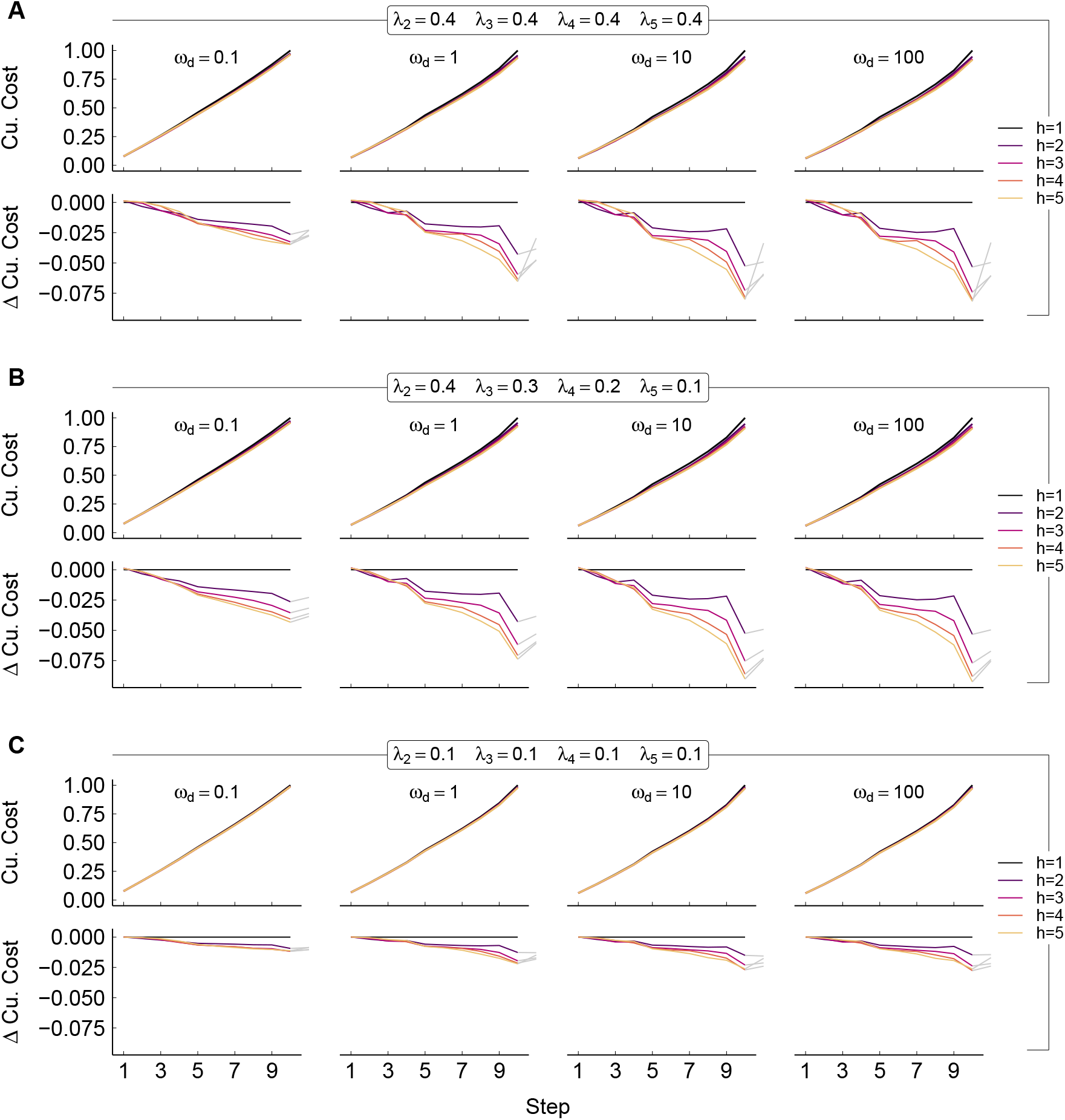
Simulated cumulative cost under the planning model, with various sets of model parameters. Cumulative cost is normalized to the mean final cumulative cost under the model with *h* = 1. Fixed parameters were chosen to be representative of the best fits for participant data (*γ* = 1.5, *ω*_*s*_ = 10, *β* = 1); movement time is assumed to be a linear in the distance (*t* = 0.2 + 0.02*d*). **A:** Top row shows cumulative cost, bottom row shows the difference in cumulative cost relative to *h* = 1 up to the 10th step (colored lines) and one additional step (gray lines). **B**,**C:** Same as A, but with different values for *λ*_2_, …, *λ*_*h*_.

**Figure S2.**
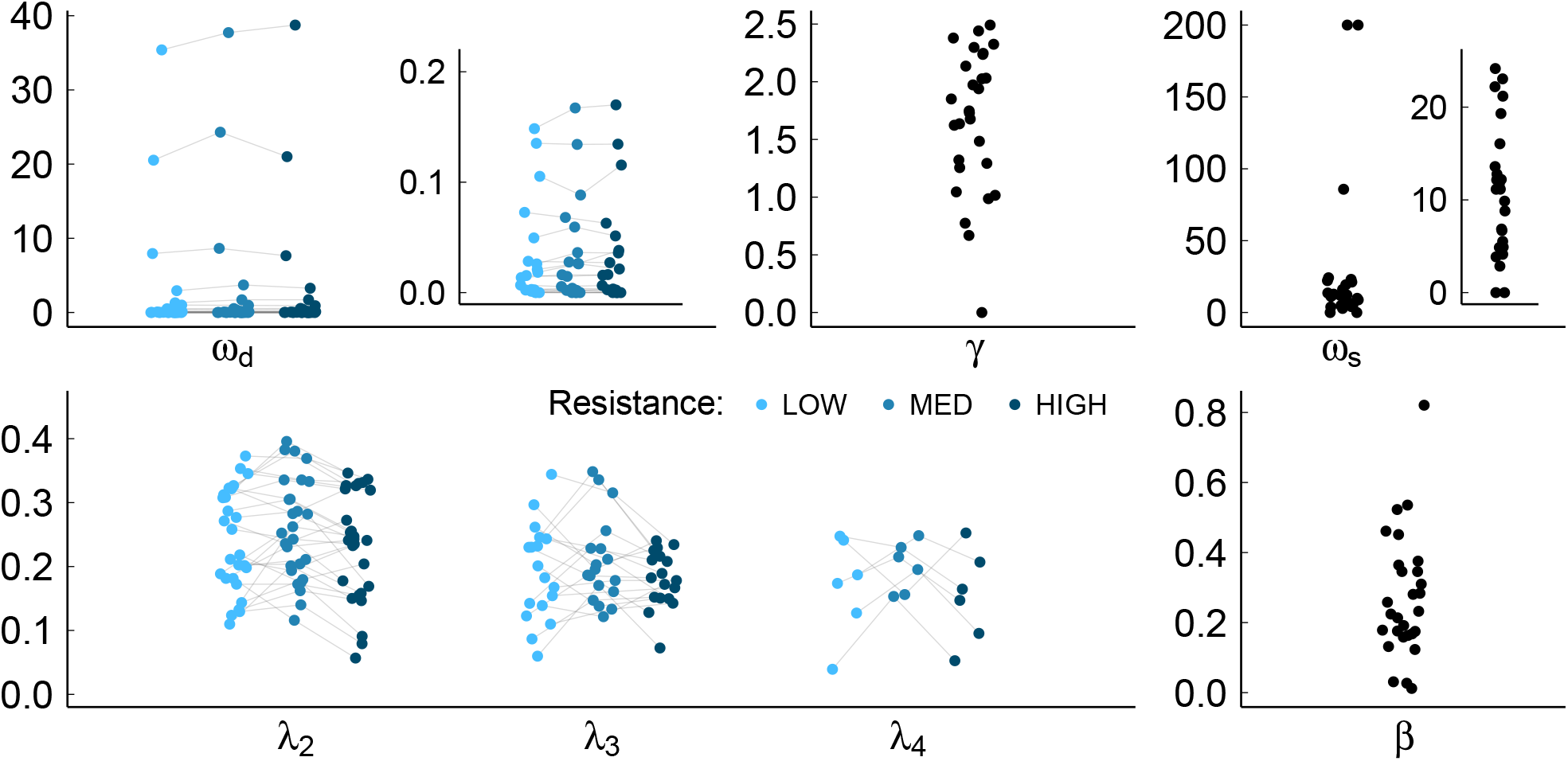
Parameters from the model best explaining all the data, for each participant.

**Table S1.** Statistical contrasts between weights for future targets 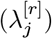 according to their index (*j* = 2, 3, 4), and according to low, medium and high resistance (*r* = *r*_1_, *r*_2_, *r*_3_). All parameters are taken from the model best explaining data across all resistance levels.

| Contrast | $\Delta$ | adjusted<br>$p$-value | 95%<br>Lower | 95%<br>Upper |
| --- | --- | --- | --- | --- |
| $\lambda_3 - \lambda_2$ | -0.083 | <0.001 | -0.102 | -0.064 |
| $\lambda_4 - \lambda_2$ | -0.118 | <0.001 | -0.148 | -0.088 |
| $\lambda_4 - \lambda_3$ | -0.035 | 0.019 | -0.065 | -0.005 |
| $\lambda^{[r_2]} - \lambda^{[r_1]}$ | 0.019 | 0.056 | -0.0004 | 0.039 |
| $\lambda^{[r_3]} - \lambda^{[r_1]}$ | -0.010 | 0.457 | -0.030 | 0.010 |
| $\lambda^{[r_3]} - \lambda^{[r_2]}$ | -0.029 | 0.002 | -0.049 | -0.010 |

**Table S2.** Statistical contrasts between weights for future targets 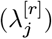 according to their index (*j* = 2, 3, 4, 5), and according to low, medium and high resistance (*r* = *r*_1_, *r*_2_, *r*_3_). All parameters are taken from the model best explaining data for the corresponding resistance level only.

| Contrast | $\Delta$ | adjusted<br>$p$-value | 95%<br>Lower | 95%<br>Upper |
| --- | --- | --- | --- | --- |
| $\lambda_3 - \lambda_2$ | -0.076 | <0.001 | -0.100 | -0.051 |
| $\lambda_4 - \lambda_2$ | -0.112 | <0.001 | -0.144 | -0.080 |
| $\lambda_4 - \lambda_3$ | -0.037 | 0.020 | -0.069 | -0.004 |
| $\lambda_5 - \lambda_2$ | -0.005 | 0.999 | -0.148 | 0.137 |
| $\lambda_5 - \lambda_3$ | 0.070 | 0.578 | -0.073 | 0.213 |
| $\lambda_5 - \lambda_4$ | 0.107 | 0.220 | -0.037 | 0.251 |
| $\lambda^{[r_2]} - \lambda^{[r_1]}$ | 0.008 | 0.737 | -0.017 | 0.032 |
| $\lambda^{[r_3]} - \lambda^{[r_1]}$ | 0.000 | 0.999 | -0.024 | 0.025 |
| $\lambda^{[r_3]} - \lambda^{[r_2]}$ | -0.007 | 0.739 | -0.031 | 0.016 |

## References

Ariani G, Diedrichsen J. Sequence learning is driven by improvements in motor planning. Journal of Neurophysiology 121: 2088–2100, 2019.

Ariani G, Kordjazi N, Pruszynski JA, Diedrichsen J. The Planning Horizon for Movement Sequences. eNeuro 8, 2021.

Ariani G, Shahbazi M, Diedrichsen J. Cortical Areas for Planning Sequences before and during Movement. Journal of Neuroscience 45, 2025.

Assarioti EE, Beers RJ van, Smeets JBJ, Wijk BCM van. Reaching Distance Influences Perceptual Decisions. European Journal of Neuroscience 61: e70006, 2025.

Bashford L, Kobak D, Diedrichsen J, Mehring C. Motor skill learning decreases movement variability and increases planning horizon. Journal of Neurophysiology 127: 995–1006, 2022.

Berlot E, Popp NJ, Grafton ST, Diedrichsen J. Combining Repetition Suppression and Pattern Analysis Provides New Insights into the Role of M1 and Parietal Areas in Skilled Sequential Actions. Journal of Neuroscience 41: 7649–7661, 2021.

Burk D, Ingram JN, Franklin DW, Shadlen MN, Wolpert DM. Motor Effort Alters Changes of Mind in Sensorimotor Decision Making. PLOS ONE 9: e92681, 2014.

Byrd RH, Lu P, Nocedal J, Zhu C. A Limited Memory Algorithm for Bound Constrained Optimization. SIAM Journal on Scientific Computing 16: 1190–1208, 1995.

Carroll TJ, McNamee D, Ingram JN, Wolpert DM. Rapid Visuomotor Responses Reflect Value-Based Decisions. The Journal of Neuroscience 39: 3906–3920, 2019.

Christopoulos V, Schrater PR. Dynamic Integration of Value Information into a Common Probability Currency as a Theory for Flexible Decision Making. PLoS Computational Biology 11: e1004402, 2015.

De Comite A, Lefèvre P, Crevecoeur F. Continuous evaluation of cost-to-go for flexible reaching control and online decisions. PLOS Computational Biology 19: e1011493, 2023.

Diamond JS, Wolpert DM, Flanagan JR. Rapid target foraging with reach or gaze: The hand looks further ahead than the eye. PLOS Computational Biology 13: e1005504, 2017.

Dolstra E. The purely functional software deployment model [Online]. 2006. https://dspace.library.uu.nl/handle/1874/7540 [4 Aug. 2025].

Eluchans M, Maselli A, Lancia GL, Pezzulo G. Eye and hand coarticulation during problem solving reveals hierarchically organized planning. bioRxiv 2024.

Fitts PM. The information capacity of the human motor system in controlling the amplitude of movement. Journal of Experimental Psychology 47: 381–391, 1954.

Gallivan JP, Johnsrude IS, Randall Flanagan J. Planning Ahead: Object-Directed Sequential Actions Decoded from Human Frontoparietal and Occipitotemporal Networks. Cerebral Cortex 26: 708–730, 2016.

Hart PE, Nilsson NJ, Raphael B. A Formal Basis for the Heuristic Determination of Minimum Cost Paths. IEEE Transactions on Systems Science and Cybernetics 4: 100–107, 1968.

Howard IS, Ingram JN, Wolpert DM. A modular planar robotic manipulandum with end-point torque control. Journal of Neuroscience Methods 181: 199–211, 2009.

Kalidindi HT, Crevecoeur F. Task-dependent coarticulation of movement sequences. eLife 13: RP96854, 2024.

Kashefi M, Diedrichsen J, Pruszynski JA. Motor Sequence Learning Involves Better Prediction of the Next Action and Optimization of Movement Trajectories. Journal of Neuroscience 45, 2025.

Kashefi M, Reschechtko S, Ariani G, Shahbazi M, Tan A, Diedrichsen J, Pruszynski JA. Future movement plans interact in sequential arm movements. eLife 13, 2024.

Kistemaker DA, Wong JD, Gribble PL. The Central Nervous System Does Not Minimize Energy Cost in Arm Movements. Journal of Neurophysiology 104: 2985–2994, 2010.

Kistemaker DA, Wong JD, Gribble PL. The cost of moving optimally: Kinematic path selection. Journal of Neurophysiology 112: 1815–1824, 2014.

Landau WM, Levin MG, Furneaux B, Company EL and. Crew. cluster: Crew Launcher Plugins for Traditional High-Performance Computing Clusters [Online]. 2025a. https://cran.r-project.org/web/packages/crew.cluster/index.html [30 Jun. 2025].

Landau WM, Warkentin MT, Edmondson M, Oliver S, Mahr T, Company EL and. Targets: Dynamic Function-Oriented ‘Make’-Like Declarative Pipelines [Online]. 2025b. https://cran.r-project.org/web/packages/targets/index.html [30 Jun. 2025].

Landau WM, Woodie D, Company EL and. Crew: A Distributed Worker Launcher Framework [Online]. 2025c. https://cran.r-project.org/web/packages/crew/index.html [30 Jun. 2025].

Manzone JX, Welsh TN. Explicit effort may not influence perceptuomotor decisionmaking. Experimental Brain Research 241: 2715–2733, 2023.

Mattar MG, Lengyel M. Planning in the brain. Neuron 110: 914–934, 2022.

Mistry M, Theodorou E, Schaal S, Kawato M. Optimal control of reaching includes kinematic constraints. Journal of Neurophysiology 110: 1–11, 2013.

Nashed JY, Crevecoeur F, Scott SH. Rapid Online Selection between Multiple Motor Plans. The Journal of Neuroscience 34: 1769–1780, 2014.

Pierrieau E, Lepage J-F, Bernier P-M. Action Costs Rapidly and Automatically Interfere with Reward-Based Decision-Making in a Reaching Task. eNeuro 8: ENEURO.0247–21.2021, 2021.

R Core Team. R: A language and environment for statistical computing [Online]. R Foundation for Statistical Computing 2025. https://www.R-project.org/.

Summerside EM, Shadmehr R, Ahmed AA. Vigor of reaching movements: Reward discounts the cost of effort. Journal of Neurophysiology 119: 2347–2357, 2018.

The MathWorks Inc. MATLAB version: 23.2 (R2023b) [Online]. 2023. https://www.mathworks.com.

Ushey K, Wickham H. Renv: Project Environments [Online]. 2025. https://cran.r-project.org/web/packages/renv/index.html [30 Jun. 2025].

Velázquez-Vargas CA, Taylor JA. Learning to Move and Plan like the Knight: Sequential Decision Making with a Novel Motor Mapping. Computational Brain & Behavior, 2025. doi:10.1007/s42113-025-00245-9.

Yokoi A, Diedrichsen J. Neural Organization of Hierarchical Motor Sequence Representations in the Human Neocortex. Neuron 103: 1178–1190.e7, 2019.

Zimnik AJ, Churchland MM. Independent generation of sequence elements by motor cortex. Nature Neuroscience 24: 412–424, 2021.

